# CTD-dependent and - independent mechanisms govern co-transcriptional capping of Pol II transcripts

**DOI:** 10.1101/269977

**Authors:** Melvin Noe Gonzalez, Shigeo Sato, Chieri Tomomori-Sato, Joan W. Conaway, Ronald C. Conaway

## Abstract

Co-transcriptional capping of RNA polymerase II (Pol II) transcripts by capping enzyme proceeds orders of magnitude more efficiently than capping of free RNA. Previous studies brought to light a role for the phosphorylated Pol II CTD in activation of co-transcriptional capping; however, CTD phosphorylation alone could not account for the observed magnitude of activation. Here, we exploit a defined Pol II transcription system that supports both CTD phosphorylation and robust activation of capping to dissect the mechanism of co-transcriptional capping. Taken together, our findings identify a novel CTD-independent, but Pol II-mediated, mechanism that functions in parallel with CTD-dependent processes to ensure optimal capping, and they support a “tethering” model for the mechanism of activation.

## Introduction

Messenger RNA and other transcripts synthesized by RNA polymerase II (Pol II) are distinguished by the presence of a 5’ guanosine cap. The cap is added by the capping enzyme to nascent Pol II transcripts bearing 5’-triphosphate ends, where it aids in subsequent steps of RNA maturation, transport, translation, and other processes(Ramanathan et al., 2016; Topisirovic et al., 2011). A remarkable property of the capping reaction is its selectivity for Pol II transcripts. Despite the presence in cells of abundant Pol I and Pol III transcripts with 5’ triphosphate ends, Pol II transcripts are the primary targets for capping by the capping enzyme(Ghosh and Lima, 2010; Ramanathan et al., 2016). Consequently, how this exquisite selectivity is accomplished has been of major interest.

An important clue to the selectivity of capping enzyme came from the discovery that co-transcriptional capping of Pol II transcripts is substantially more efficient than capping of free RNA; indeed, the specific activity of capping enzyme for nascent transcripts emerging from elongating Pol II is several orders of magnitude greater than its specific activity for free RNA (Moteki and Price, 2002). This revelation argued that inherent features of the Pol II transcription complex are responsible for dramatically activating capping and, in so doing, ensuring selectivity of capping enzyme.

Though it is presently not known exactly why co-transcriptional capping is so efficient, previous studies have implicated the phosphorylated Pol II CTD in both recruitment and activation of the capping enzyme (Cho et al., 1997; McCracken et al., 1997). Pol II is distinguished from Pol I and Pol III by the presence of a unique carboxyl-terminal domain (CTD) on its largest subunit, RPB1. The Pol II CTD, conserved from yeast to humans, consists of a tandemly repeated heptapeptide motif with consensus sequence Y_1_S_2_P_3_T_4_S_5_P_6_S_7_, which is subject to extensive phosphorylation.

Additional studies have shed considerable light on biochemical mechanisms underlying activation of co-transcriptional capping of Pol II transcripts, leading to the formulation of several non-mutually exclusive models for how the phosphorylated CTD might activate capping. One model proposes that the phosphorylated CTD activates capping by recruiting and tethering the capping enzyme to elongating Pol II in the vicinity of the emerging nascent transcript (tethering model). This model is supported by evidence that the capping enzyme binds specifically and stably to GST-CTD or CTD heptapeptides phosphorylated at either serine 2 (pSer2) or serine 5 (pSer5) (Ho and Shuman, 1999; Ho et al., 1998; McCracken et al., 1997; Rodriguez et al., 2000). A second model proposes that the pSer5-CTD activates capping by allosterically activating capping enzyme (allosteric activation model). This model is supported by evidence that binding of capping enzyme to CTD heptapeptide repeats phosphorylated on serine 5, but not on serine 2, increases formation of the covalent capping enzyme-GMP complex, an intermediate during transfer of the 5’-guanosine cap to Pol II transcripts (Cho et al., 1998; Ho and Shuman, 1999).

Despite this evidence, the relative importance of CTD phosphorylation for activation of capping has been questioned, since blocking CTD phosphorylation only partially inhibits co-transcriptional capping. CDK7, the protein kinase associated with the Pol II initiation factor TFIIH, preferentially phosphorylates CTD Ser5 and Ser7 (Compe and Egly, 2016). Addition of CDK7 inhibitors to block CTD phosphorylation in transcription complexes assembled in nuclear extracts led to only a modest, ~4-fold reduction in capping efficiency (Moteki and Price, 2002; Nilson et al., 2015), suggesting that other activation mechanisms likely contribute to the activation of RNA capping.

In this report, we exploit a defined Pol II transcription system that supports both CTD phosphorylation and robust activation of co-transcriptional capping to dissect the mechanism of capping. As described below, our findings define a CTD-independent mechanism that functions in parallel with CTD-dependent processes to ensure maximal capping. In addition, we report mechanistic experiments that argue that a combination of CTD-independent and CTD-dependent tethering mechanisms likely play a dominant role in activation of co-transcriptional capping.

## Methods

### Materials

Unlabeled ultrapure ribonucleoside 5’ triphosphates were from GE Healthcare, 3’-O-Methyl Guanosine-5’ Triphosphate (3’-OMeGTP, cat. no. TM03-002) was from Ribomed, and [α-32P] CTP, GTP, or UTP (all 3000 Ci/mmol) were from Perkin Elmer. Rnasin Plus RNase Inhibitor (40 units/μl, cat. no. N2611) was from Promega. Bovine Serum Albumin (20 mg/ml, cat. no. B9000S), 2x RNA Loading Dye (cat. no. B0363S), and Yeast Inorganic Pyrophosphatase (100 units/ml, cat. no. NEBM2403S) were from New England Biolabs. RNase T1 (1000 units/μl, cat. no. EN0541), GlycoBlue Coprecipitant (15mg/ml, cat. no. AM9516), and Proteinase K Solution (20 mg/ml, cat. no. 25530049) were from Life Technologies Invitrogen. Protease Inhibitor for mammalian cell extracts (cat. no. P8340) and Protease Inhibitor Cocktail for His-Tag purifications (cat. no. P8849) were from Sigma, and 10 mM THZ1 Hydrochloride in DMSO (cat. no. HY-80013A) was obtained from MedChem Express.

RNA 5’ Polyphosphatase (cat. no. RP8092H), Terminator 5’-Phosphate-Dependent Exonuclease (cat. no. TER51020), and Tobacco Acid Pyrophosphatase (TAP; cat. no. T81050, discontinued) were from Epicentre, and Decapping Pyrophosphohydrolase (DppH; cat. no. 003436004, discontinued) was from Tebu-bio.

Magnetic beads coupled to streptavidin (Dynabeads MyOne Streptavidin C1, Dynabeads MyOne Strepatividin T1, or Dynabeads M-280) were from Life Technologies Invitrogen. Anti-FLAG M2 affinity gel (cat. no. A2220) and FLAG peptide (cat. no. F3290) were from Sigma. MaXtract High Density tubes (1.5 ml, 129046) from Qiagen.

DNA oligonucleotides were obtained from IDT (see primer table for purity specifications). 5’-triphosphorylated RNAs (containing ~15% un-modified RNA) were obtained from Trilink.

Anti-Rpb1-NTD Rabbit mAb (D8L4Y) was from Cell Signaling; anti-Rpb1 antibody N-20 (sc-899) and anti-Rpb2 antibody E-12 (sc-166803) were from Santa Cruz; Rat anti-RNA Pol II CTD phospho Ser5 monoclonal antibody (cat. no. 61085) was from Active Motif. IRDye 800CW Goat Anti-Rat IgG (925-32219) was from LiCor; Donkey anti-Rabbit IgG Alexa Fluor 680 (A10043) and Donkey anti-Mouse IgG Alexa Fluor 680 (A10038) were from Invitrogen.

### Preparation of RNA Polymerase II and Transcription Factors

RNA Polymerase II and TFIIH were purified as described from rat liver nuclear extracts (Conaway et al., 1996). Recombinant yeast TBP (Conaway et al., 1991), recombinant rat TFIIB (Tsuboi et al., 1992). Recombinant human TFIIE was prepared as described (Peterson et al., 1991), except that the 56-kDA subunit was expressed in E. coli strain BL21(DE3)-pLysS. TFIIF RAP30 and RAP74 subunits were amplified from human cDNA and inserted into pETDuet-1 vector (Novagen) MCS1 (His-tag) and MCS2 (no tag), respectively. Intact TFIIF was expressed in E. coli strain BL21(DE3)-RIL and purified on Ni-NTA agarose. Purified S. pombe Pol II was a gift from Henrik Spähr and was purified as described (Banks et al., 2007).

### Plasmids and Immobilized Templates

Plasmid pMLT-Gal4(5)-G219, made in a pGEM3 backbone, contains five 17-bp Gal4-binding sites (each separated by 2 bp) 14 bp upstream of the AdML promoter from - 50 to +10, followed by a 219 bp G-less cassette. pMLT-Gal4(5)-INS20 is identical pMLT-Gal4(5)-G219 except for an insertion of “GGG” after position +20 relative to the AdML transcription start site.

The 861 bp biotinylated G23 DNA template was prepared by PCR using pMLT-Gal4(5)-INS20 as template. The 5’-primer (pMLTG5_FOR_Biotin) was biotinylated at its 5’ end and was complementary to the sequence 204 bp upstream of the first Gal4-binding site; the 3’ primer (pMLTG5_REV) was complementary to the sequence 197 bp downstream of the second G-less cassette. PCR products were purified using a QIAquick/MinElute PCR purification Kit (Qiagen).

The 99 bp biotinylated G21 DNA template contained AdML promoter sequences - 36 to +10 and included a 20 bp G-less cassette. G21 template was prepared by annealing 1 nmole of non-template strand, 5’-biotinylated DNA oligo to 2 nmole of template strand DNA oligo in 50μl of 50 mM KCl, 5 mM MgCl_2_, 25mM Tris-Cl pH 7.5. The oligo mixture was incubated for 5 min at 95 °C, then slow cooled over 72 min at 1 °C per min in a PCR machine. Biotinylated G21 DNA template was stable for at least 2 months at 4 °C.

To prepare immobilized templates, ~ 1 mg of M-280, MyOne C1, or MyOne T1 magnetic beads (Invitrogen) was incubated with ~15-50 pmole (10-30 μg) of biotinylated G23 template or with 1 nmole of G21 template for 30 min at room temperature in 5 mM Tris-Cl pH 7.5, .5 mM EDTA, 1 M NaCl, collected using a Dynamag-2 magnet (ThermoFisher), and then washed three times in the same buffer. Beads were then washed an additional three times in 20 mM HEPES-NaOH pH 7.9, 20% Glycerol, 100 mM KCl, 1mM EDTA, 0.5 mg/ml bovine serum albumin, and finally resuspended in the same buffer to a final bead concentration of 10 μg/μl. Templates immobilized on magnetic beads were kept at 4 °C and were stable for at least 6 months.

Yeast vector p-YN132 containing full-length mouse capping enzyme was a generous gift from Stewart Shuman. cDNA encoding capping enzyme was released from this plasmid by digestion with Nde1 and Xho1 and subcloned into pET15b, which encodes an in-frame N-terminal 6x HisTag.

### Expression and purification of mammalian capping enzyme

6x His-mammalian capping enzyme was expressed in E. coli strain BL21(DE3)-RIL and purified on Ni-NTA agarose. Capping enzyme was dialyzed for 2 hours in 10K MWCO Slide-A-Lyzer cassettes (Thermo-Fisher) against 50 mM Tris-Cl pH 8, 50 mM NaCl, 2 mM EDTA, 10 % glycerol, 5 mM sodium pyrophosphate (to de-guanylate capping enzyme), and then dialyzed overnight against the same buffer lacking sodium pyrophosphate. The purity and concentration of capping enzyme was assessed on Coomassie Blue-stained SDS gels, using BSA as a standard. Purified enzyme was aliquoted and kept at −80 °C.

### Promoter-dependent Transcription

Unless otherwise mentioned in the figure legend, PICs were assembled for 30 min at 30 °C in 30 or 60 μl reaction mixtures containing 50-100 ng of G21 or G23 template immobilized on magnetic beads, ~10 ng of recombinant TFIIB, ~400 ng of recombinant TFIIF, ~20 ng of recombinant TFIIE, ~300 ng of TFIIH, 50 ng of yeast TBP and 0.02 units of RNA polymerase II in 3 mM HEPES-NaOH pH 7.9, 20 mM Tris-HCl pH 7.9, 60 mM KCl, 0.5 mM DTT, 0.5 mg/ml bovine serum albumin, 2% polyvinyl alcohol, 3% glycerol, 8 mM MgCl_2_ (<u>B</u>ase <u>T</u>ranscription <u>B</u>uffer or BTB), supplemented with 20 U of RNasin Plus (Promega N2611).

To synthesize 21mers, PICs assembled on G21 templates were incubated with 100 μM (Figs 2D and S1) or 125 μM (Figs 1C and 2A) 3’OMeGTP, 50 μM ATP, 50 μM UTP, 2 μM CTP, 10 μCi α-^32^P-CTP (3000 Ci/mmol), 8 mM MgCl_2_, and 1 μl of T1 RNase for the indicated times.

**Figure 1.**
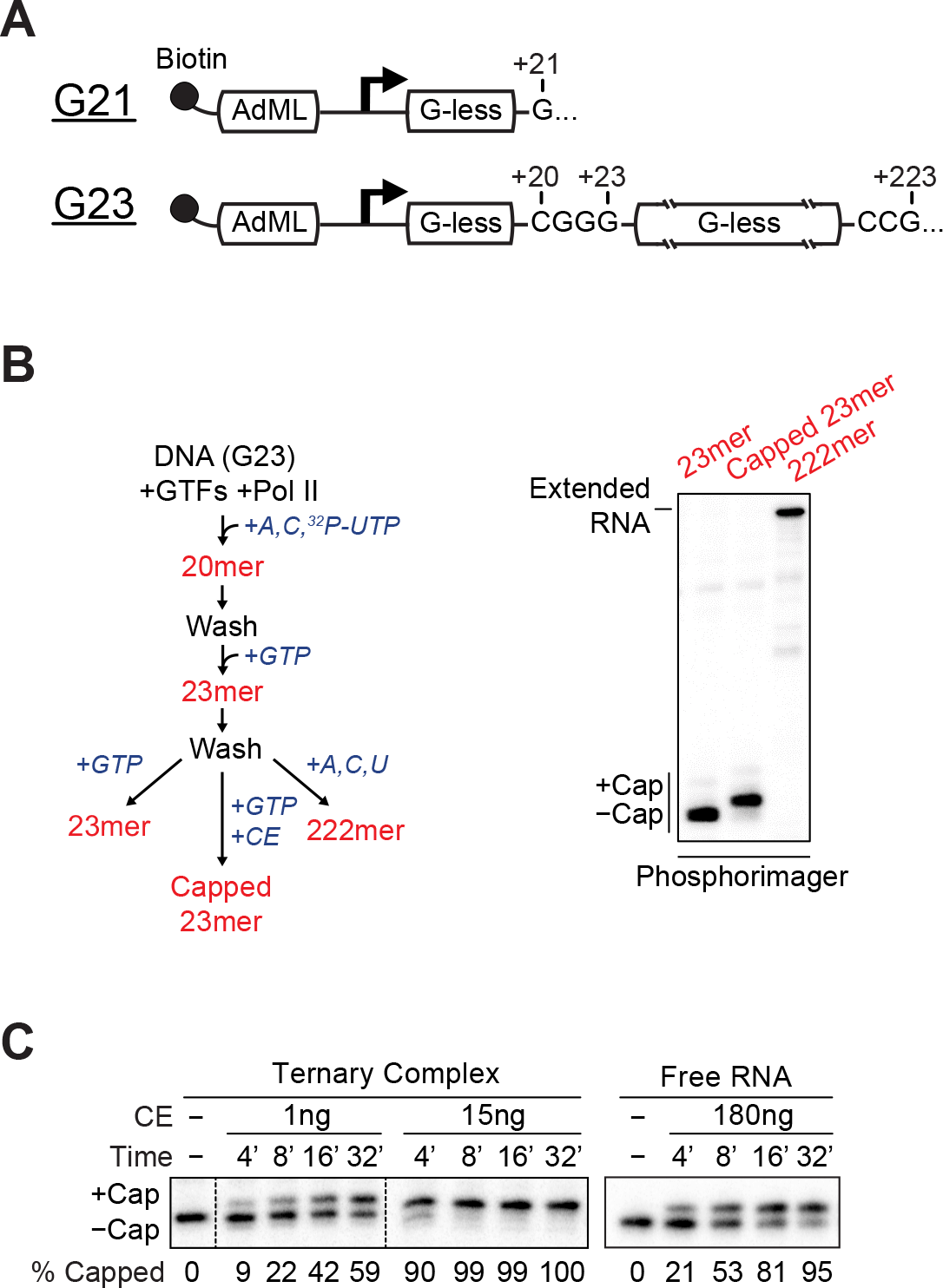
Co-transcriptional capping activation in a defined enzyme system. ***A,*** Biotinylated DNA templates used for promoter-dependent transcription. Both contain the Adenovirus 2 Major Late core promoter (AdML) followed by one (G21) or two (G23) G-less cassettes. ***B,*** 23mer transcripts in washed ternary complexes were prepared according to the diagram and incubated with GTP (lane 1), GTP and 5 ng of capping enzyme (CE) (lane 2), or ATP, CTP, and UTP (lane 3). In this and subsequent figures, radiolabeled transcripts were resolved by denaturing gel electrophoresis and detected using a phosphorimager. ***C,*** Kinetics of co-transcriptional capping and capping of free RNA. Free RNA or washed ternary complexes containing 21mers were incubated for varying lengths of time with 50 μM GTP and the indicated amounts of capping enzyme (CE).

**Figure 2.**
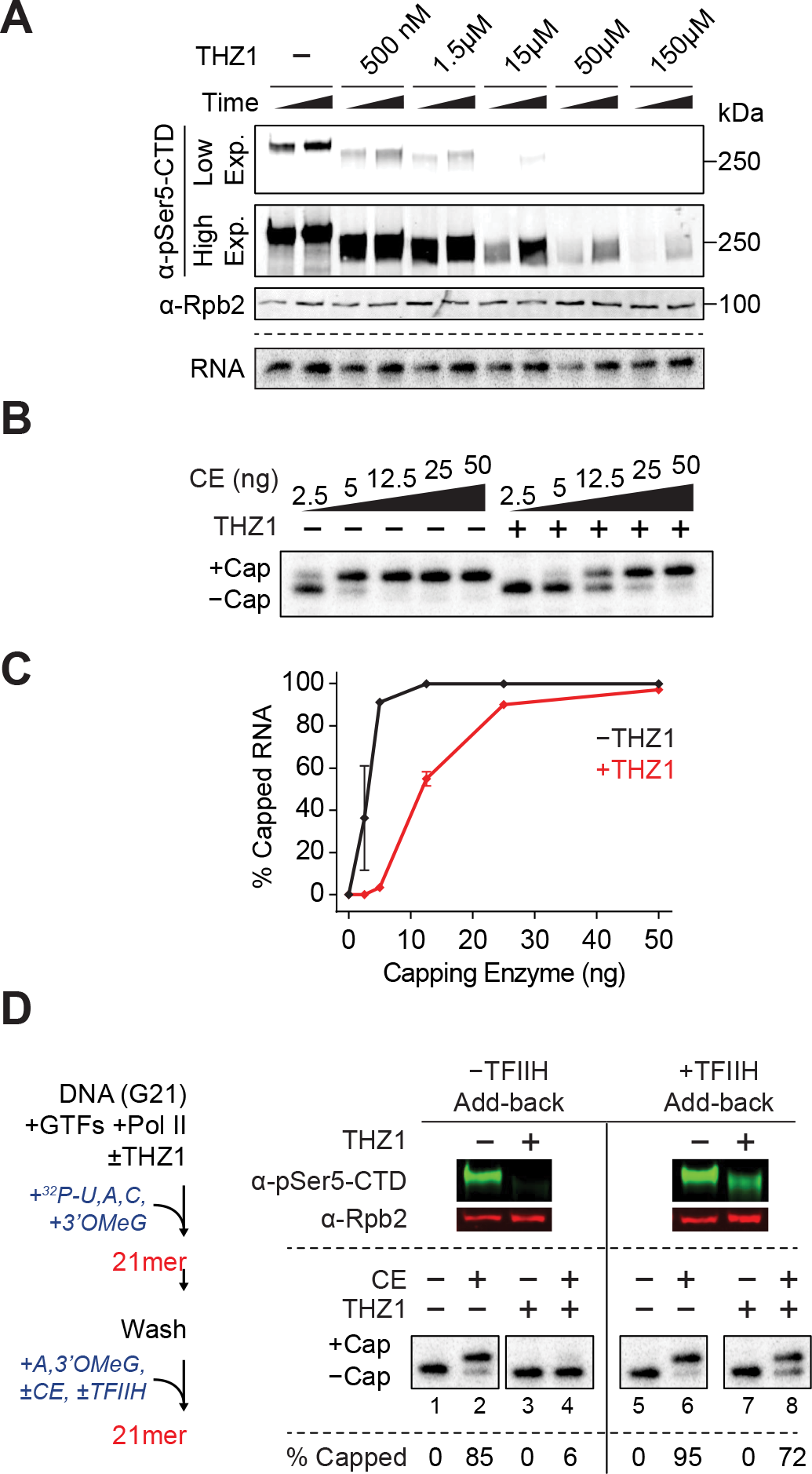
Cdk7 strongly stimulates co-transcriptional capping in the purified enzyme system. ***A,*** 21mers were synthesized in parallel reactions with unlabeled (upper panel) or radiolabeled (lower panel) ribonucleoside triphosphates in the presence of DMSO (-) or increasing amounts of THZ1; reactions were stopped after 15 or 60 min. Upper panel, reaction products were analyzed by western blotting using antibodies against Ser5 phosphorylated Rpb1 (α-pSer5-CTD); two different exposures of the same image are shown. As a control for equal loading of Pol II in each lane, the same blot was probed with antibodies against Rpb2. Lower panel, radiolabeled transcripts were analyzed on denaturing gels and detected by phosphorimaging. ***B,*** Washed transcription complexes containing 23mers synthesized with or without 150 μM THZ1 were incubated for 4 min with GTP and increasing amounts of capping enzyme (CE). ***C,*** Graph shows mean and range of 2 independent reactions performed as in ***B***. ***D,*** As diagrammed on the left, washed transcription complexes containing 21mers were prepared in the presence of DMSO or 100-150 μM THZ1 and incubated for 15 min with 50 μM ATP and 100 μM 3’OMeG, with or without 3 ng of capping enzyme. For +TFIIH add-back reactions (lanes 5-8), 300 ng of purified TFIIH was added with capping enzyme. Reactions were assayed by Western blotting for Pol II CTD phosphorylation status (top) or for RNA capping (bottom). % Capped indicates average of two independent reactions.

**Table 1.**
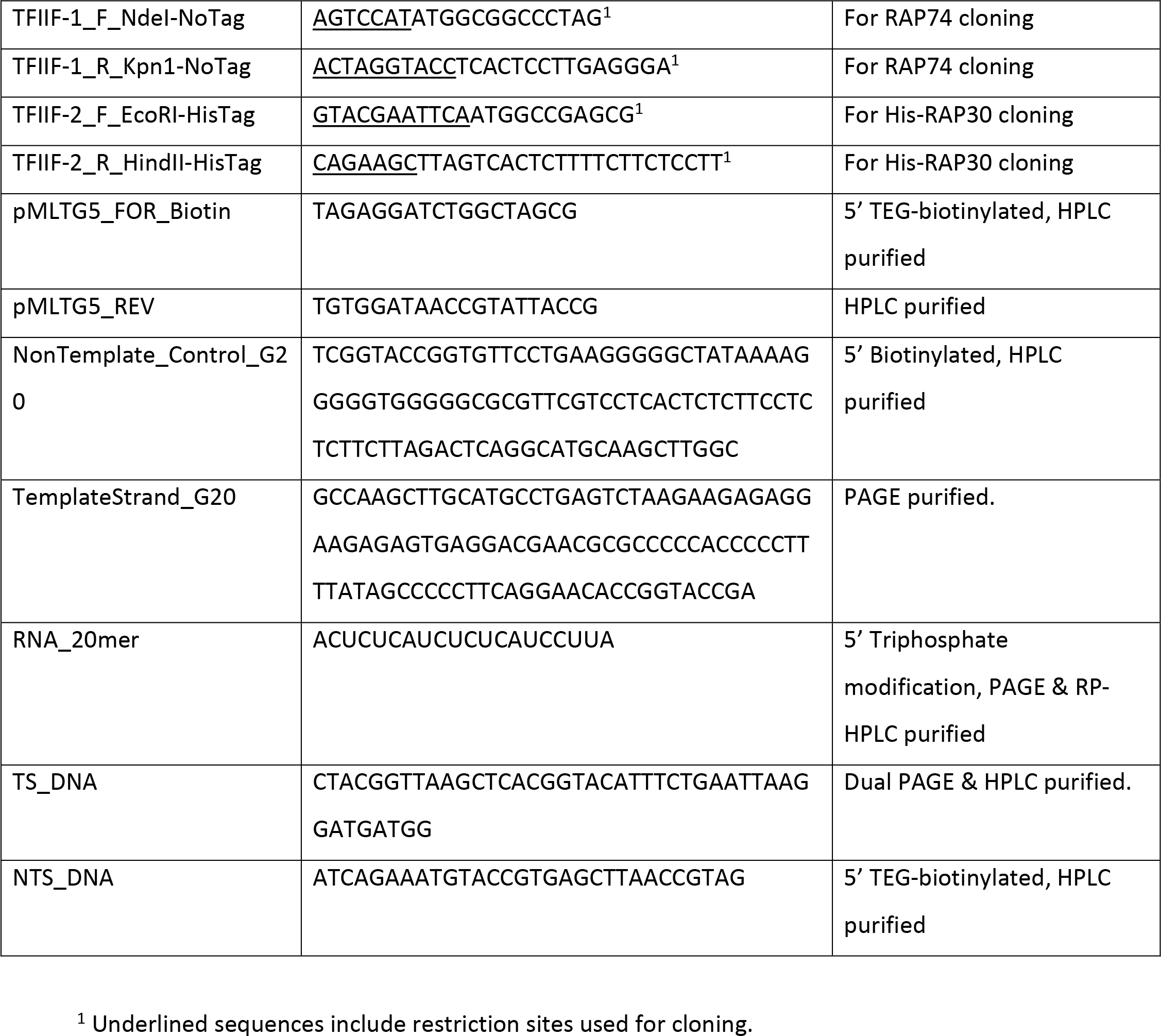
Primers and Oligos.

To synthesize 23mers, 222mers, and 223mers, PICs were assembled on G23 templates immobilized on magnetic beads, and Pol II was walked to various positions by successive incubations at 30°C with appropriate combinations of nucleotides. Transcription was initiated by addition of 25 μM ATP, 5 μM CTP, 5 μM UTP, 10 μCi of α-^32^P-UTP (3000 Ci/mmol). After 5 min, reactions were supplemented with an additional 50 μM ATP, 50 μM CTP, and 50 μM UTP and incubated an additional 2 min to generate 20 mers. Immobilized transcription complexes were collected and washed twice with a volume of high salt wash buffer (HSB, containing 3 mM HEPES-NaOH pH 7.9, 20 mM Tris-HCl pH 7.9, 1 M KCl, 0.5 mM DTT, 0.5 mg/ml bovine serum albumin, 0.2% polyvinyl alcohol, and 3% glycerol) equivalent to the initial reaction volume, followed by two more washes in low salt wash buffer (LSB, containing 3 mM HEPES-NaOH pH 7.9, 20 mM Tris-HCl pH 7.9, 60 mM KCl, 0.5 mM DTT, 0.5 mg/ml bovine serum albumin, 0.2% polyvinyl alcohol, and 3% glycerol). Transcription complexes were then incubated for 2 min in BTB containing 5 μM GTP to generate 23mers, washed twice with HSB and twice with LSB. When necessary, transcription complexes containing 23mers were resuspended in BTB containing 500 μM ATP, 500 μM CTP, and 500 μM UTP, incubated for 30 min to allow synthesis of 222mers and washed as described above. Washed transcription complexes containing 23mers or 222mers were used as substrates for RNA capping reactions or analyzed by denaturing polyacrylamide gel electrophoresis. 222 nt transcripts were extended to 223mers during capping reactions, since GTP used as capping substrate allowed for addition of one nucleotide to transcripts.

Where indicated, 150 μM THZ1 was included during PIC assembly and synthesis of 20mers or 21mers; to control for solvent effects an equivalent volume of DMSO was included in control reactions for these experiments. All subsequent steps, except for washes, included 50 μM THZ1, even in control reactions, to normalize for any post-initiation effect(s) of THZ1.

### Artificial Pol II Elongation Complexes

For assembly of artificial elongation complexes, 1 nmole of non-template DNA was immobilized on magnetic beads and washed as described above. Immobilized oligo was stable for at least 6 months at 4 °C. To begin assembly of artificial elongation complexes, 20 pmol of RNA 20-mer oligo with a 5’ triphosphate (RNA_20mer) were annealed to 10 pmol of template strand DNA oligo (TS_DNA) in 10μl 25 mM Tris pH 7.5, 50 mM KCl, 5 mM MgCl_2_. Reactions were incubated 5 min at 45 °C, then incubated for 12 cycles of 2 min each, starting at 43 °C and decreasing the temperature 2 °C per cycle in a PCR machine. All further incubations were at 30 °C. 1 pmol of template strand:RNA hybrid was incubated with 0.02 units of Pol II in 50 mM Tris pH 7.5, 50 mM KCl, 5 mM MgCl_2_, 2% polyvinyl alcohol, 3% Glycerol, 0.5 mg/ml BSA, 0.5 mM DTT for 10 min. An equal volume of the same buffer supplemented with 5 pmol of non-template strand DNA oligo (NTS_DNA) immobilized on magnetic beads was then added to the reaction and incubated for 10 min at 37 °C.Immobilized scaffolds were then washed by collecting samples in magnet for 2 min, then resuspended in an equal volume of LSB. To generate transcription complexes containing radiolabeled 23mers, immobilized scaffolds were collected, resuspended in 25 μl of BTB containing 0.6 μM ATP and 10 μCi of α-^32^P-UTP (3000 Ci/mmol), and incubated for 10 min. Following this incubation, 5μl of BTB supplemented with 5 μM ATP and 5 μM UTP was added, reactions were incubated a further 5 min, and the resulting transcription complexes were washed twice with 3 mM HEPES-NaOH pH 7.9, 20 mM Tris-HCl pH 7.9, 60 mM KCl, 0.5 mM DTT, 0.5 mg/ml bovine serum albumin, 0.2% polyvinyl alcohol, 3% glycerol.

Washed transcription complexes were processed differently depending on the nature of the experiment. To walk Pol II along the template to generate transcripts of the desired lengths, transcription complexes were resuspended in BTB supplemented with the appropriate combinations of 20 μM NTPs and incubated for 10 min. For phosphorylation by TFIIH, transcription complexes were resuspended in BTB supplemented with 50 μm ATP and 1 μl (~300 ng) of purified TFIIH for 10 min. For capping, washed complexes were processed as described below. In multi-step reactions, such as that of figure 3B, transcription complexes were washed twice with 3 mM HEPES-NaOH pH 7.9, 20 mM Tris-HCl pH 7.9, 60 mM KCl, .5 mM DTT, 0.5 mg/ml bovine serum albumin, 0.2% polyvinyl alcohol, 3% glycerol between steps.

**Figure 3.**
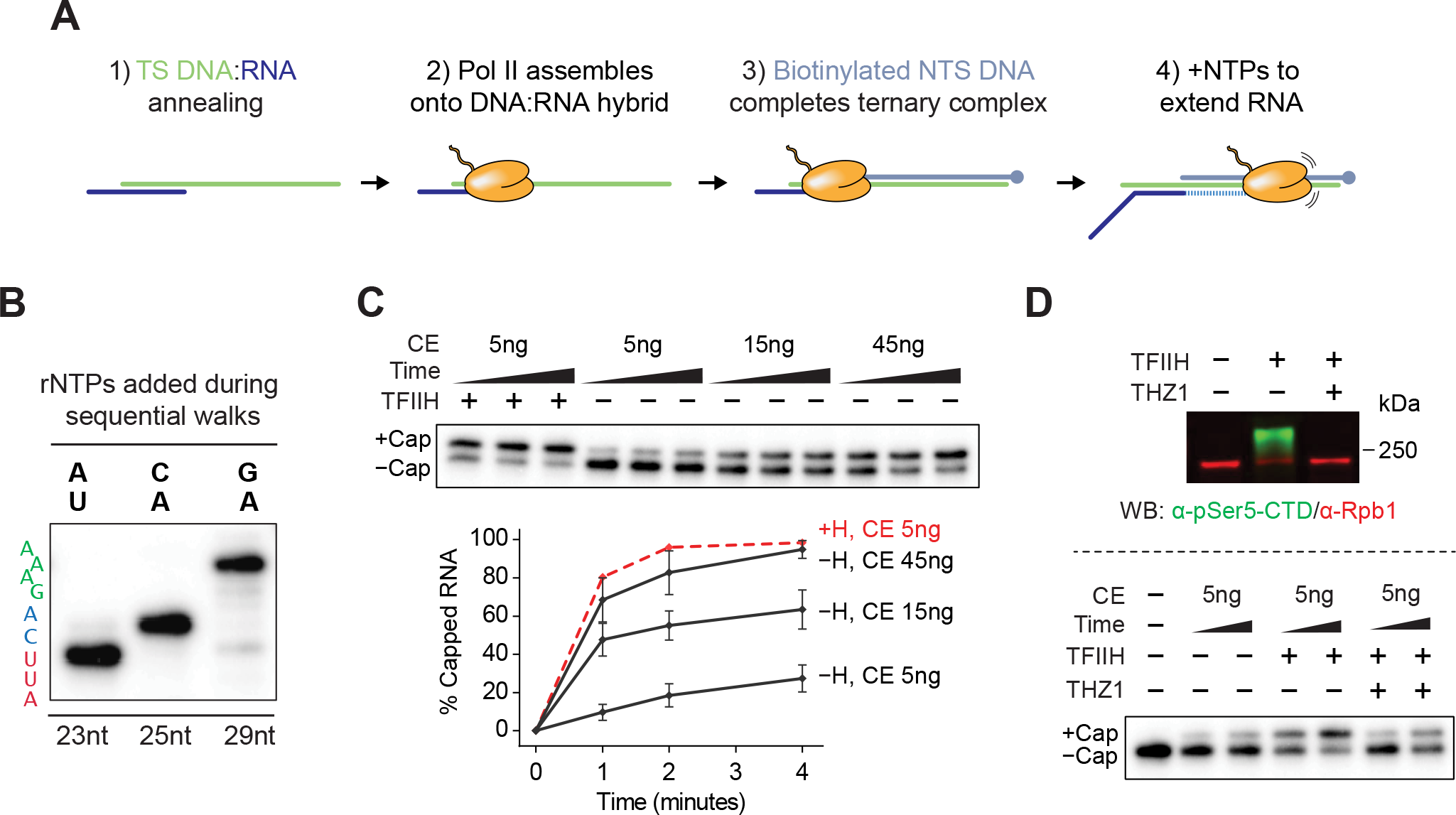
TFIIH-dependent activation of co-transcriptional capping in artificial ternary complexes assembled on DNA:RNA scaffolds. ***A,*** Scheme for assembly of artificial ternary elongation complexes on DNA:RNA scaffolds. See text and “Materials & Methods” for details. ***B,*** Representative gel image showing step-wise increase of RNA length following addition of appropriate combinations of rNTPs. 20-nt synthetic RNA in artificial ternary complexes was extended to position +23 with 5 μM ATP and 10 μCi α-32P UTP (first lane), washed and walked to position +25 with 20 μM CTP and ATP (second lane), and finally washed and walked to position +29 by with 20 μM ATP and GTP (last lane). Sequence on left denotes RNA sequence from +21 (bottom) to +29 (top). ***C,*** 23mers in artificial ternary complexes were incubated with buffer or 300 ng of purified TFIIH for 10 minutes, washed, and incubated for 1, 2, or 4 min with 50 μM GTP and the indicated amounts of capping enzyme. Graph shows mean and range of data from 2 independent reactions. ***D,*** Artificial ternary complexes with 23mers were pre-incubated with DMSO (-) or 150 μM THZ1 for 20 min, supplemented with buffer or 300 ng of purified TFIIH, and incubated for an additional 10 min. After washing, ternary complexes were incubated with 50 μM GTP, and with buffer (lane 1) or 5 ng of capping enzyme. Aliquots of capping reactions were stopped after 4 min and assayed by Western blot for Pol II CTD phosphorylation status (top panel) or after 1 and 4 min and assayed for RNA capping (bottom panel). Shown is a representative example of experiments performed in duplicate.

### Free RNA

Radiolabeled 23mer RNA was generated using artificial Pol II elongation complexes and purified by phenol-chloroform-isoamyl alcohol extraction, chloroform extraction, and ethanol precipitation as described below. Purified RNA was resuspended in 5 μl of H_2_O per reaction and used as a substrate for capping reactions.

### RNA Capping

Free RNA or washed transcription complexes with transcripts initiated from the promoter on the G23 template or transcripts in artificial ternary complexes were suspended in BTB supplemented with 50 μM GTP and 0.1 unit of inorganic yeast polyphosphatase and then transferred to new tubes containing the indicated amounts of capping enzyme. To assay capping of transcripts initiated from the promoter on the G21 template, capping enzyme was added to pre-assembled PICs along with nucleotides for RNA synthesis; note that these reactions included 100-125 μM 3’OMeGTP instead of GTP.

### Purification of capped and uncapped transcripts and analysis by denaturing gel electrophoresis

Transcription or capping reactions were stopped by addition of 60 μl 10 mM Tris-HCl (pH 7.5), 300 mM NaCl, 0.5 mM EDTA, 0.2% SDS and 2 μl of 20 mg/ml proteinase K, 2μl of GlycoBlue 15 mg/ml (Invitrogen AM9516), and enough H_2_O the final volume of solution to 124 μl, including the volume of the original reaction mix. Samples were then extracted once with 124 μl of phenol:choloroform:isoamyl alcohol (25:24:1) and once with chloroform:isoamyl alcohol (24:1) using MaXtract High Density tubes (Qiagen), brought to 0.3 M sodium acetate by addition of 12.4 μl of 3 M sodium acetate pH 5.2, ethanol precipitated, and washed with 70% ethanol. After removal of the final ethanol wash, RNA pellets were air dried for 3 min, resuspended in 1X RNA Loading Dye, heated at 70 °C for 10 min, spun 4 min at 2000 x g, and resolved on a denaturing gel (15% PAGE 1:19 bis/tris, 7.0 M urea). Radiolabeled gels were exposed to a phosphorimager (Molecular Dynamics or Amersham Biosciences) and scanned using a Typhoon Trio imager (Amersham Biosciences). Images were quantified using ImageQuantTL (GE Healthcare) and plotted using Graphpad Prism (Version 6.05).

% capped RNA was determined by measuring the ratio of capped RNA signal/total RNA (capped + uncapped) and normalized to the maximum obtainable capping. Maximum capping of transcripts initiated from promoter was consistently 100%. For analysis of free RNA or artificial Pol II elongation complexes, the maximum % capped RNA was determined to be 85%, likely due to incomplete triphosphorylation of synthetic RNA.

### Western blotting

Protein samples were boiled 10 min at 70 °C, and loaded into 5% handcasted SDS-PAGE gels or commercially available pre-casted gels (Biorad 3450002). After electrophoresis, proteins were transferred to a PVDF membrane (Millipore Immobilon-FL). Membranes were blocked for 30 min using Odyssey Blocking Buffer PBS (LICOR), incubated overnight with primary antibody that had been diluted in Odyssey Blocking Buffer supplemented with 0.1% Tween-20, washed with TBS containing 0.05% Tween 20, incubated with the appropriate secondary antibodies at room temperature, washed again, and then finally scanned using a LICOR Odyssey Scanning Instrument.

### Purification of human Pol II containing RPB1 lacking the CTD

Gibson assembly was used to generate a DNA fragment encoding RPB1 lacking the CTD (RPB1-ΔCTD, amino acids 1-1592) from 3 separate DNA fragments. Fragment 1 included a 5’ XhoI site followed by nucleotides 1-1449 of NM_000937 CDS; fragment 2 included nucleotides 1425-2829, and fragment 3 included nucleotides 2808-4776 followed by stop codon and BamHI site; these were synthesized by PCR, assembled, and cloned into pcDNA5 (Life Technologies). This insert was sub-cloned into a modified pcDNA5/FRT/TO vector that encodes an in-frame N-terminal 3xFLAG tag, and co-transfected with pOG44 into Flip-In T-Rex 293 cells with FuGENE6 (Promega). Stably transfected cells were selected using 100 μg/ml of hygromycin. Hygromycin-resistant cell clones were treated with 2 μg/ml of doxycycline for 48 hrs to induce RPB1-ΔCTD, and protein expression was confirmed by immunoblot with anti-FLAG mAb.

Intact nuclei was prepared essentially as described (Aygun et al., 2008). Briefly, stable Flip-In T-Rex 293 F:RPB1-ΔCTD cells were grown to near confluence in 4 roller bottles, treated with 2 μg/ml doxycycline, and harvested 24 hrs later. Cells were lysed with 10 mM HEPES-NaOH pH 7.9, 0.34 M sucrose, 3 mM CaCl_2_, 2 mM magnesium acetate, 0.1 mM EDTA, 1 mM DTT, 0.5% Nonidet P-40, and protease inhibitors (Sigma), strained through a 70 μM filter, and pelleted by centrifugation. Nuclei were washed with the same buffer without detergent before continuing with preparation of total nuclear extracts (Dignam, Roeder) (Dignam et al., 1983). Nuclear extracts were applied to M2-agarose beads, and F:RPB1-ΔCTD protein complex was purified as previously described (Tomomori-Sato et al., 2013).

### Assay for formation of GMP-capping enzyme intermediate

Free RNA or washed artificial ternary complexes were suspended in BTB supplemented with 0.1 unit of inorganic yeast polyphosphatase and 10 μCi of α-^32^P-GTP (3000 Ci/mmol) instead of GTP, and then transferred to new tubes containing 5 ng of capping enzyme. Reactions were stopped with 1x Laemmli Sample Buffer at the indicated times, boiled for 10 min at 70 °C, and loaded into 4-20% gradient gel (Biorad). After electrophoresis, gels were fixed in 40% methanol, 10% acetic acid for 1 hr, and rinsed with water. Radiolabeled gels were exposed to a phosphorimager and processed as described above.

## Results

### Activation of Co-transcriptional Capping in a Minimal Pol II Transcription System

To begin to investigate the mechanism underlying activation of co-transcriptional capping of Pol II transcripts, we took advantage of a defined Pol II transcription system. In this system, transcription is reconstituted with purified RNA polymerase II and TFIIH, recombinant TBP, TFIIB, TFIIE, and TFIIF, and DNA templates containing the adenovirus 2 major late promoter (AdML) followed by one (G21) or two (G23) G-less cassettes (Figure 1A).

Immobilized G23 templates were incubated with Pol II and initiation factors to assemble pre-initiation complexes (PICs). Following PIC assembly, ATP, α-^32^P UTP, and CTP were added to allow synthesis of 20 nucleotide (nt) transcripts and then washed to remove unincorporated rNTPs. 20mers were walked to 23mers by addition of GTP and washed again. The resulting ternary transcription complexes were incubated with or without recombinant mammalian capping enzyme and with GTP, which serves as the GMP donor for the capping reaction. Capping of nascent transcripts was monitored by a commonly used assay that detects an electrophoretic mobility shift of approximately 1 nt in capped transcripts (Chiu et al., 2002; Mandal et al., 2004), indicating addition of a 5′ cap to the RNA (Figure 1B, compare lanes 1 and 2). Confirming that the 23mers used as substrates for RNA capping were associated with transcribing Pol II, they could be chased quantitatively into longer transcripts upon addition of ATP, CTP, and UTP to allow transcription to proceed through the second G-less cassette in the G23 template (Figure 1B, third lane). Enzymatic digestion of these RNA products using a combination of cap-sensitive phosphatases and exonucleases confirmed that the shift in mobility was due to capping of the nascent transcript (Figure S1). Furthermore, and consistent with previous findings (Moteki and Price, 2002), we observe that the specific activity of capping enzyme for transcripts associated with the Pol II transcription complex is substantially greater than it is for free RNA. As shown in Figure 1C, 15 ng of capping enzyme was sufficient to cap 90% of transcripts in co-transcriptional capping reactions in 4 min, while it took more than 30 min to achieve a similar amount of capping of free RNA with 180 ng of capping enzyme (Figure 1C).

### The TFIIH associated kinase activates co-transcriptional capping in the minimal Pol II transcriptional system

Because previous studies implicated phosphorylation of the Pol II CTD as a key step in activation of capping, we first explored the contribution of CTD phosphorylation to capping in our minimal Pol II transcription system. The TFIIH associated CDK7 kinase is solely responsible for CTD phosphorylation in our system.

THZ1 has been shown to act as a covalent inhibitor of the TFIIH-associated CDK7 kinase and of CDK7-dependent serine 5 phosphorylation on the Rpb1 CTD *in vitro* and *in vivo* (Kwiatkowski et al., 2014). Preinitiation complexes were assembled on immobilized templates, and G-less transcripts were synthesized in the presence and absence of THZ1. Addition of THZ1 decreased Ser-5 phosphorylation on the CTD (pSer5-CTD) during transcription by 90% at 1 μM and achieved near complete inhibition by 150 μM (Figure 2A). As expected from previous results demonstrating that CTD phosphorylation is not required for basal transcription with purified factors *in vitro* (Serizawa et al., 1993), THZ1 had no major effect on RNA synthesis even at the highest concentration used (Figure 2A, bottom panel).

Addition of THZ1 substantially decreased, but did not completely inhibit, co-transcriptional capping in the reconstituted enzyme system. As shown in Figures 2B and 2C, 5 ng of capping enzyme was sufficient to cap nearly all transcripts in the absence of THZ1, while ~5 times more capping enzyme was needed to achieve a similar level of capping in the presence of 150 μM THZ1. Notably, co-transcriptional capping in the presence of THZ1 was still substantially more efficient than capping of free RNA.

To ensure that the TFIIH kinase is the target of THZ1-dependent inhibition of capping, we carried out a TFIIH “add-back” experiment. Transcription complexes containing 21mers were synthesized in the presence or absence of THZ1 as diagrammed in Fig. 2D. After extensive washing to remove THZ1, capping reactions were performed with or without addition of new TFIIH, in the presence of a low concentration of capping enzyme, such that THZ1 almost completely blocks capping. As expected, Ser-5 CTD phosphorylation (Figure 2D, upper panel) and RNA capping (Figure 2D, lower panel) were inhibited when using 5 ng of capping enzyme. However, add-back of untreated TFIIH, following THZ1 inhibition, rescued Ser-5 CTD phosphorylation and RNA capping.

### The TFIIH kinase activates co-transcriptional capping of transcripts elongated by Pol II pre-assembled on synthetic DNA:RNA transcription bubbles

Thus far our results indicate that the TFIIH kinase can activate co-transcriptional capping of transcripts initiated by Pol II from a promoter in the presence of a minimal set of initiation factors. A limitation of these assays is that TFIIH is required not only for CTD phosphorylation but also for transcription initiation; moreover, using these assays we can not distinguish between the possibilities (i) that the residual amount of TFIIH kinase activity observed even in the presence of high THZ1 concentrations is sufficient to activate co-transcriptional capping or (ii) that co-transcriptional capping can be activated by both phosphorylation-dependent and - independent events.

To investigate more directly the requirement for the TFIIH kinase, we sought to simplify the transcription system further using artificial Pol II elongation complexes pre-assembled on synthetic DNA:RNA transcription bubbles, which allow transcription without a promoter and without general transcription factors. This methodology has been successfully used to study the structures and function of ternary complexes assembled with budding yeast and mammalian Pol II (e.g. (Kellinger et al., 2012; Kireeva et al., 2000; Wang et al., 2009; Wang et al., 2015)) and to obtain an EM structure of fission yeast Pol II bound to capping enzyme (Martinez-Rucobo et al., 2015). As discussed below, we found that activation of co-transcriptional capping of transcripts associated with artificial Pol II elongation complexes was as robust as activation in the minimal Pol II transcription system.

DNA template strand (TS) and 20 nt RNA oligonucleotides with 5’-triphosphate ends were annealed to form a DNA:RNA hybrid, incubated with purified mammalian Pol II to correctly position the enzyme on the scaffold, and then supplemented with a molar excess of biotinylated DNA non-template strand (NTS) oligo to close the ternary complex (Figure 3A). Immobilized ternary complexes were then washed extensively to remove unbound Pol II and DNA and RNA oligos, followed by addition of appropriate combinations of rNTPs to walk the RNA to the desired length (Fig. 3B).

Co-transcriptional capping in ternary elongation complexes assembled on DNA:RNA scaffolds recapitulated features of capping of promoter-specific transcripts in our defined enzyme system. Although TFIIH was not essential for co-transcriptional capping in these ternary complexes, including it in reactions increased the specific activity of capping enzyme by ~10-fold (Figure 3C and Figure S2). Moreover, THZ1 dramatically inhibited formation of p-Ser5 CTD (Figure 3D, upper panel) and reduced co-transcriptional RNA capping in reactions containing TFIIH (Figure 3D, lower panel), indicating that the catalytic activity of CDK7 is needed for TFIIH-dependent activation of capping in ternary elongation complexes. Thus, TFIIH-dependent activation of capping does not require initiation from a promoter and is independent of any of the other initiation factors.

### Direct evidence that the Pol II CTD is the target of the TFIIH kinase

Our finding that Pol II elongation complexes assembled on DNA:RNA scaffolds faithfully recapitulate co-transcriptional RNA capping in the absence of a promoter and general transcription factors argues (i) that TFIIH-dependent capping activation depends solely on features of the Pol II elongation complex and (ii) that Pol II is the sole target of protein kinase activity required for activation of capping.

To test directly whether the Pol II CTD is the target of the TFIIH kinase, we asked whether TFIIH stimulates capping in ternary elongation complexes assembled with mutant Pol II lacking the CTD. To accomplish this, we used FLAG-immunopurification to prepare CTD-deficient Pol II from a 293 cell line expressing an Rpb1 mutant lacking the entire CTD and containing an N-terminal FLAG epitope tag (Figure 4A).

**Figure 4.**
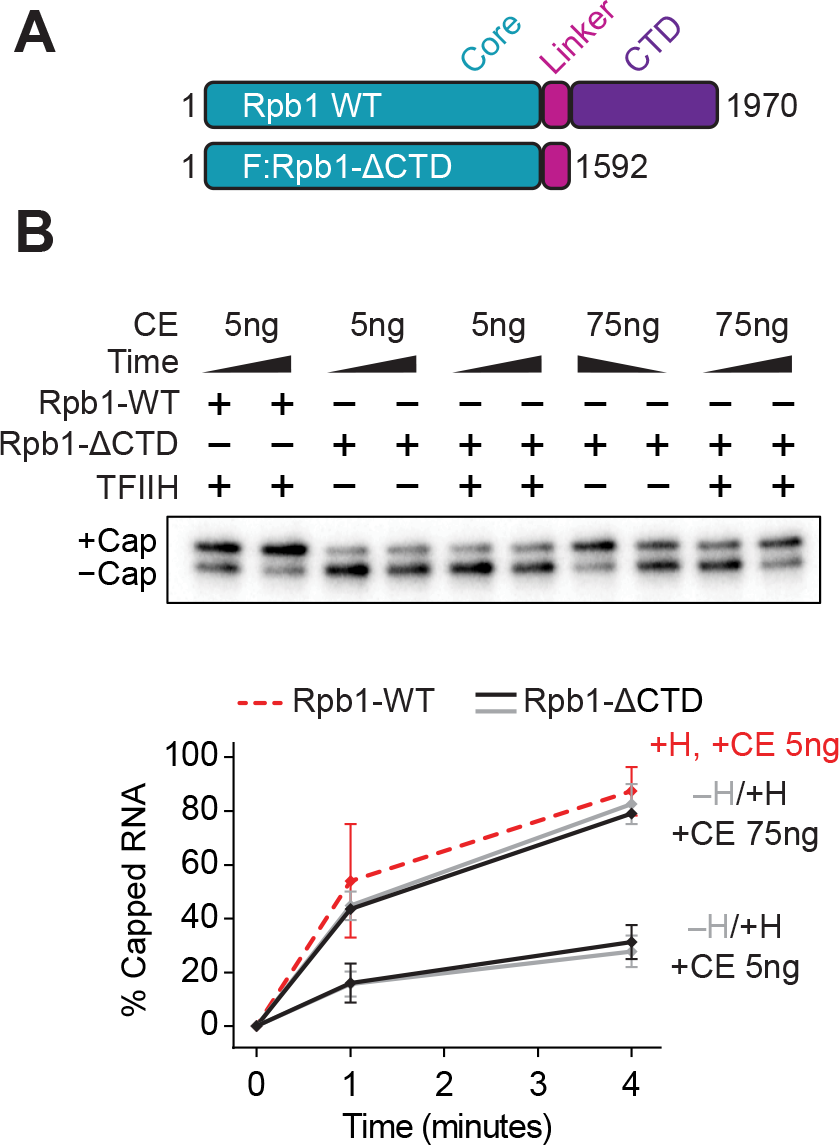
Pol II CTD is necessary for TFIIH-dependent activation of co-transcriptional capping. ***A,*** Diagram of wild type (WT) Rpb1 and FLAG-tagged Rpb1 mutant lacking the CTD (F:Rpb1-ΔCTD). Numbers denote amino acid positions. ***B,*** Artificial ternary complexes containing 23mers were assembled with WT RNA Pol II (Rpb1-WT) or RNA Pol II lacking the CTD (Rpb1-ΔCTD), incubated with buffer or 300 ng of purified TFIIH for 10 min, washed, and then incubated for 1 or 4 min with 50 μM GTP and 5 ng or 75 ng of capping enzyme. 5μl and 30μl of total 30μl reaction were loaded for WT and ΔCTD Pol II respectively. Graph shows mean and range of 2 independent reactions.

In the absence of TFIIH, there was no major difference between the efficiency of capping in ternary elongation complexes containing Pol II with or without the CTD (compare Figures 3C and 4B). Whereas TFIIH strongly stimulated capping in ternary complexes containing wild type Pol II, it had no effect on either the rate or extent of capping when added to ternary complexes assembled with CTD-deficient Pol II (Fig. 4B), indicating that an intact CTD is the direct target for the TFIIH-dependent activation of co-transcriptional capping. Nevertheless, co-transcriptional capping in the absence of TFIIH and of the CTD (Fig. 4B) remains much more efficient than capping of free RNA (Figs. 1C), arguing that additional features of the ternary elongation complex, independent of the CTD, also contribute to capping activation. This observation is consistent with a prior report that capping of RNA in transcription complexes that had been assembled in nuclear extracts, washed with high salt, and treated with chymotrypsin to remove the CTD is more efficient than capping of free RNA (Moteki and Price, 2002). These prior studies did not, however, rule out the possibilities (i) that factor(s) other than the ternary complex remain after high salt washes and enhance capping or (ii) that one or a few CTD repeats remain after proteolysis.

### Species-specific interactions of capping enzyme with the body of Pol II support co-transcriptional capping

To explore the nature of CTD-independent activation of capping, we considered the possibility that proper presentation of the 5’ triphosphate ends of transcripts emerging from the Pol II exit channel is important for CTD-independent activation of capping. Alternatively, capping enzyme could have additional Pol II binding sites outside of the CTD. Indeed, structural studies have provided evidence for contacts between yeast capping enzyme and surfaces on the body of yeast Pol II, either in the multihelical “foot” domain of Rpb1 (Suh et al., 2010) or near the RNA exit channel (Martinez-Rucobo et al., 2015). Such contacts might enhance capping by positioning the capping enzyme so that it can capture the 5’ end of the nascent transcript as it emerges from the RNA exit channel; however, the contribution of Pol II body-capping enzyme interactions to co-transcriptional capping has not been explored.

If CTD-independent capping activation depends solely on the conformation of the nascent transcript as it emerges from the Pol II exit channel, one would expect that capping by mammalian capping enzyme would be insensitive to the source of Pol II used to assemble ternary elongation complexes. In contrast, if co-transcriptional capping depends on contacts between capping enzyme and surfaces in the body of Pol II, maximal CTD-independent capping by mammalian capping enzyme might be achieved only with elongation complexes containing its cognate Pol II.

To address these possibilities, we assayed capping by mammalian capping enzyme using artificial elongation complexes assembled with either mammalian Pol II or Pol II from the fission yeast *Schizosaccharomyces pombe*, which is evolutionarily distant. In the absence of TFIIH-dependent CTD phosphorylation, capping in ternary complexes containing fission yeast Pol II was reduced to a much greater extent than in ternary complexes containing mammalian Pol II. In particular, about 10 times more capping enzyme was needed to cap 50% of transcripts in fission yeast Pol II ternary complexes than to cap the same fraction of transcripts in mammalian ternary complexes (Figure 5C, compare orange lines at 50%). Notably, under conditions that support complete capping of transcripts in mammalian Pol II ternary complexes (Figure 3C, 4 min reactions, ~45 ng of capping enzyme), there was almost no capping of transcripts in ternary complexes with fission yeast Pol II (Figure 5A, B). These observations argue that contacts between capping enzyme and the body of Pol II contribute to co-transcriptional capping.

**Figure 5.**
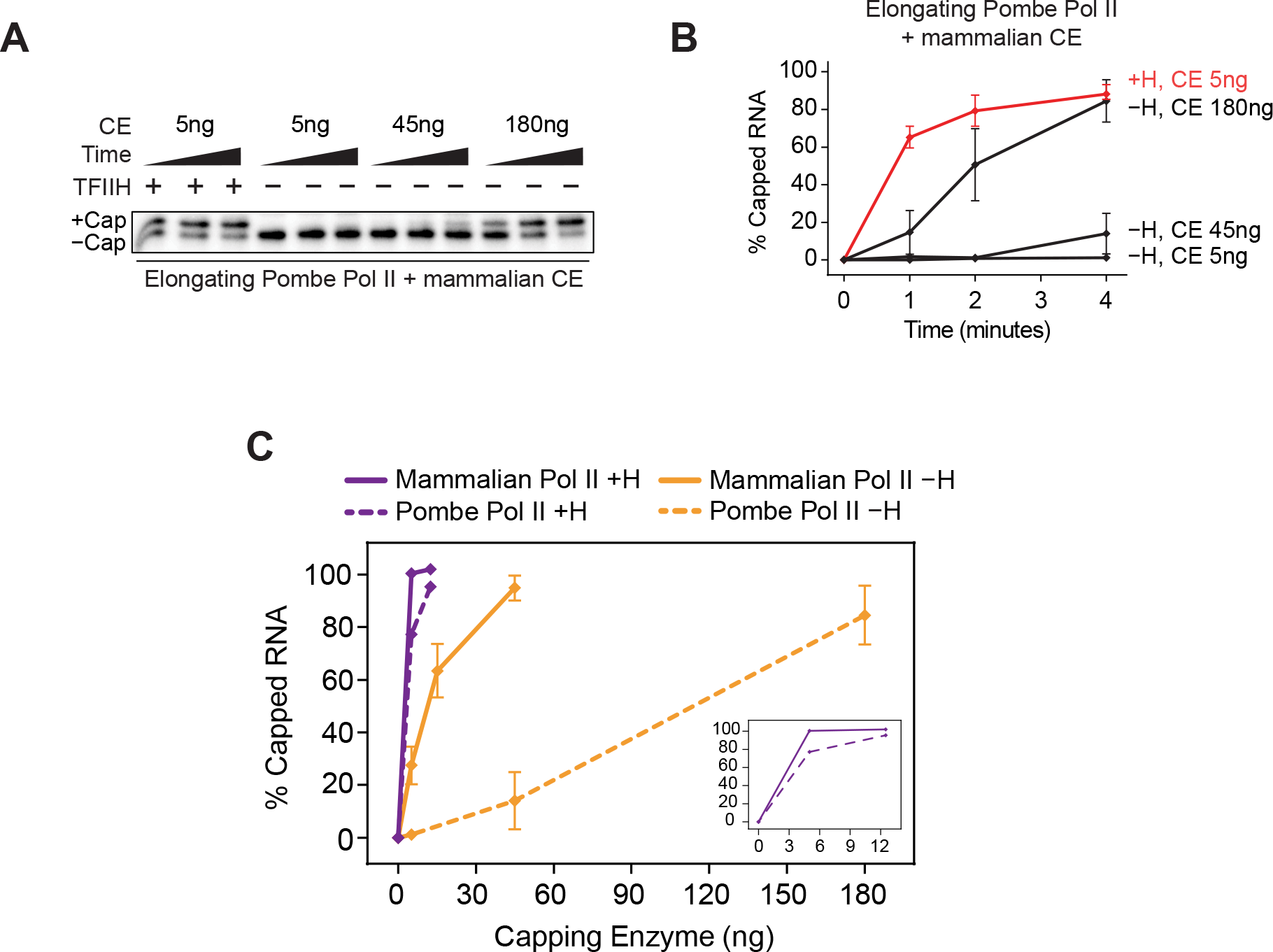
Species-specific interactions between mammalian capping enzyme and artificial ternary complexes assembled with S. Pombe Pol II. ***A,*** Artificial ternary complexes were assembled with *S. pombe* RNA Pol II, following the same protocol used for mammalian Pol II. Washed complexes containing 23mers were then incubated with buffer or 300 ng of purified TFIIH for 10 min, washed, and incubated with 50 μM GTP and capping enzyme for 1, 2, or 4 min. ***B,*** Graph shows mean and range of 2 independent reactions performed as in ***A***. ***C,*** Artificial ternary complexes containing mammalian Pol II (solid lines) or *S. pombe* Pol II (dotted lines) were incubated for 4 min with various concentrations of capping enzyme, with (purple) or without (orange) TFIIH. Graph shows mean and range from 2 independent reactions. Inset shows only reactions with TFIIH.

Interestingly, when reactions were carried out in the presence of TFIIH to phosphorylate the Pol II CTD, capping was similarly efficient in ternary complexes containing either mammalian or fission yeast Pol II (Figure 5C, compare purple lines). Our observation that TFIIH-dependent phosphorylation of the *S. pombe* Pol II CTD can restore activation of capping by mammalian capping enzyme to levels seen with mammalian Pol II suggests crosstalk between the CTD-dependent and CTD-independent mechanisms. Taken together, our findings are consistent with the model that evolutionarily conserved interaction of capping enzyme with the Pol II CTD, as well as species-specific interaction of capping enzyme with Pol II surface(s) outside the CTD contribute to activation of capping.

### A tethering model can account for activation of capping

Thus far our results argue for the existence of both CTD-dependent and CTD-independent mechanisms for activation of co-transcriptional capping of Pol II transcripts. Our results are consistent with the possibility that interaction(s) of capping enzyme with Pol II surfaces including the CTD and other sites are the major determinants of activation of capping, but they do not shed light on how these Pol II surfaces activate capping.

To begin to address this question, we carried out a systematic series of mechanistic experiments designed to explore two current, non-mutually exclusive activation models, which we refer to as the “tethering” and “allosteric activation” models (Cho et al., 1998; Ho and Shuman, 1999; Ho et al., 1998; McCracken et al., 1997; Rodriguez et al., 2000). The tethering model argues that the Pol II elongation complex acts as a scaffold to bring the capping enzyme and the 5’-triphosphate end of the nascent transcript into close proximity, thus effectively increasing the local concentrations of capping enzyme and transcript. This model does not require that the intrinsic catalytic activity of capping must be increased to account for activation of capping. The allosteric activation model argues that interaction of capping enzyme with a site or sites on Pol II is required to increase the specific activity of capping enzyme.

A key distinction between these activation models is their prediction for the fate of transcripts added in *trans*. The tethering model requires that both the capping enzyme and the nascent transcript be bound to the same Pol II scaffold in order for activation of capping to occur. Thus, free RNA added in *trans* to ternary elongation complexes would be capped as inefficiently as free RNA alone. In contrast, the allosteric activation model requires simply that capping enzyme be bound to a site or sites on Pol II to increase its intrinsic catalytic activity and predicts that activation of capping should occur with transcripts associated with Pol II or added in *trans*. We therefore asked whether transcripts added in *trans* to Pol II elongation complexes were capped with similar or different efficiencies.

First, we estimated the relative specific activities of capping enzyme for free RNA alone or free RNA mixed in *trans* with artificial Pol II elongation complexes. As seen in Figure 6A, at saturating concentrations of capping enzyme (180 and 540 ng of capping enzyme), free RNA capping required reaction times ~4-8 times longer than needed to obtain a similar level of capping in ternary elongation complexes with just 5 ng of capping enzyme. In the experiment of Figure 6B, free 29 nt RNA was added in *trans* to artificial elongation complexes containing 23 nt nascent transcripts. As expected, TFIIH strongly activated co-transcriptional capping of 23mers associated with Pol II elongation complexes. In contrast, free 29mers added in *trans* to Pol II elongation complexes were capped as inefficiently as free RNA alone, in either the presence or absence of TFIIH. These results indicate that the specific activity of capping enzyme for free RNA was not affected by the presence of either TFIIH or a phosphorylated Pol II elongation complex.

**Figure 6.**
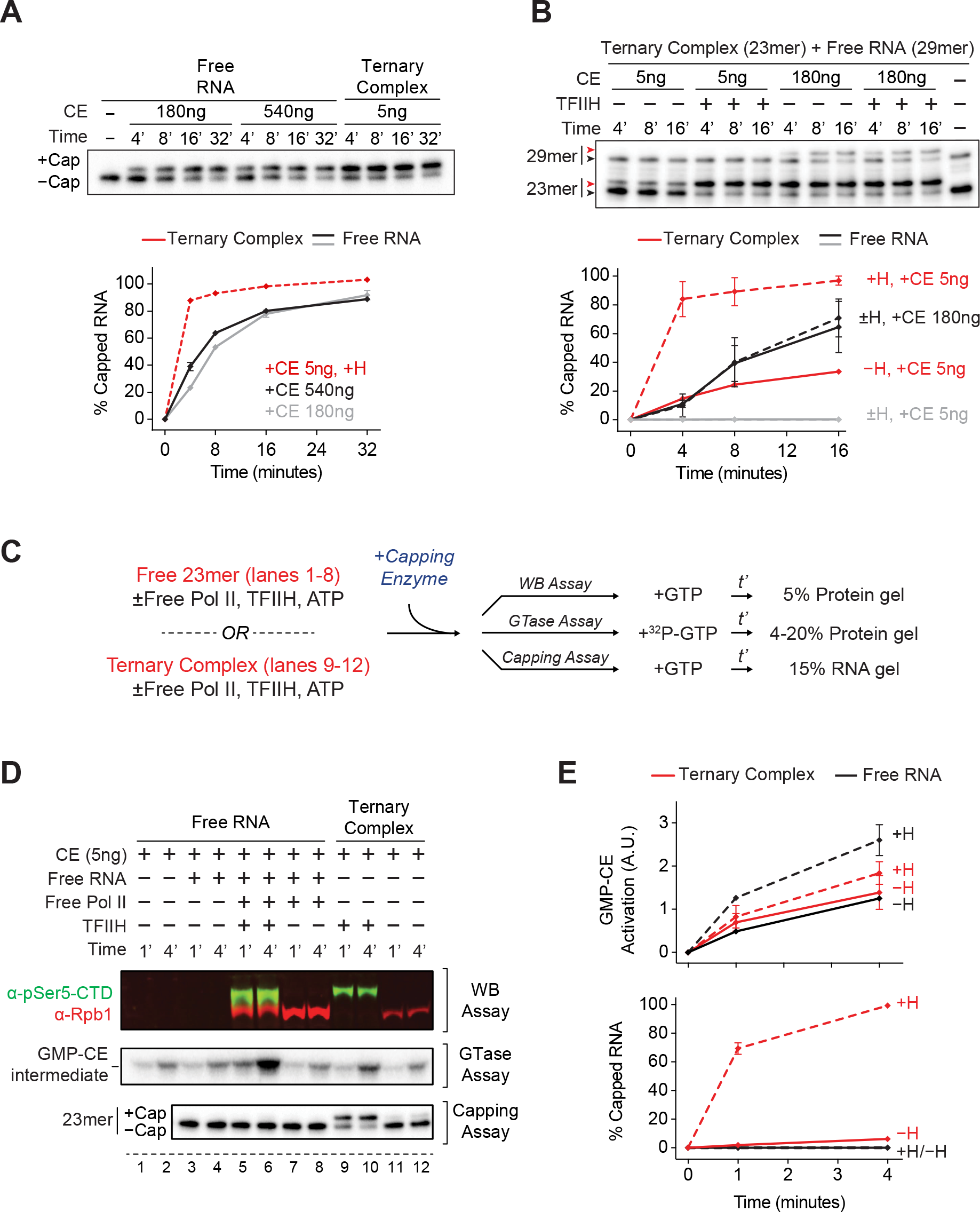
Phosphorylated Pol II provided in trans does not activate capping of free RNA. ***A,*** Kinetics of capping of free 23mer RNA or 23mers in phosphorylated artificial ternary complexes. Reactions contained 50 μM GTP and the indicated amounts of capping enzyme. The first 5 lanes are the same as those used in Fig 1C, Free RNA. Graph shows values for capping in ternary complexes from the single experiment shown in the figure and the mean and range from 2 independent experiments for free RNA capping reactions. ***B,*** Kinetics of capping of free 29mer RNA and 23mers in artificial ternary complexes, in the same reactions. Ternary complexes containing 23mers were incubated with or without 300 ng of purified TFIIH, washed, and resuspended in buffer containing 50 μM GTP. 29mer RNA was added in trans to these reactions, mixed, and reaction mixtures were transferred to new tubes containing capping enzyme. Red and black arrowheads represent capped and uncapped RNA, respectively. Graph shows mean and range of data from 2 independent reactions. ***C,*** Protocol used in assays shown in panels ***D*** and ***E***. Capping enzyme was mixed with buffer; with free 23mer RNA, in the presence or absence of pre-phosphorylated Pol II; or with 23mer RNA in phosphorylated artificial ternary complexes. Reactions were supplemented with 50 μM unlabeled GTP (WB and capping assays) or 0.3 μM α-32P GTP (guanylyltransferase assays). t’, time. ***D,*** Analysis of the reactions prepared in ***C*** by Western blotting using antibodies against Rpb1 (α-Rpb1) or against Ser5-phosphorylated Rpb1 CTD (α-pSer5-CTD), formation of α-32P GMP-capping enzyme intermediate, or RNA capping. ***E,*** Quantification of GMP-capping enzyme intermediate formation (top) or total RNA capped (bottom) from ***D***. Graphs show mean and range of 2 independent replicas of assays shown in lanes 5-12.

Evidence for the allosteric activation model comes from the finding that binding of capping enzyme to phosphorylated CTD heptapeptide repeats increases formation of the covalent GMP-capping enzyme intermediate (Cho et al., 1998; Ho and Shuman, 1999). Mammalian capping enzyme is a bifunctional enzyme possessing both RNA 5’-triphosphatase and guanylyltransferase (GTase) activities. In the first step of capping the triphosphatase hydrolyzes the RNA 5’-triphosphate to produce a diphosphate. GTP is then “loaded” into the GTase catalytic center and hydrolyzed to GMP, forming a GMP-capping enzyme intermediate. Finally, GTase transfers GMP to the 5’ diphosphate end of the RNA to form the cap (Ramanathan et al., 2016; Shuman and Hurwitz, 1981).

To test directly the correlation between allosteric activation of GTase and capping, we performed capping reactions and measured reaction products using three different assays that quantified Pol II phosphorylation (WB assay), GMP-capping enzyme intermediate (GTase assay), or RNA capping (capping assay) (Fig. 6C). Consistent with previous findings, we observed a modest increase in formation of the GMP-capping enzyme intermediate in the presence of Pol II with Ser5 phosphorylated CTD; however this increase in intermediate did not correlate with an increase in the efficiency of free RNA capping. In particular, addition of CTD phosphorylated, but not unphosphorylated, Pol II *in trans* led to an approximately 2-fold increase in formation of GMP-capping enzyme intermediate, even though there was no detectable capping of free RNA. Furthermore, formation of the GMP intermediate was increased less than 2-fold by CTD phosphorylation in reactions containing ternary Pol II elongation complexes, while co-transcriptional capping was greatly enhanced.

The results presented thus far indicate (i) that free RNA capping is not stimulated by free phosphorylated Pol II or by active ternary complexes provided in trans and (ii) that phospho-CTD-dependent changes in the rates of formation of GMP-capping enzyme intermediates do not correlate with changes in the rate of RNA capping. These observations argue that allosteric activation of GMP-capping enzyme formation through interaction with CTD-phosphorylated Pol II plays a relatively minor role in activation of co-transcriptional capping in our system.

The tethering model suggests that proximity of the RNA 5’-end to the Pol II elongation complex might contribute to capping activation. If this is the case, one might expect that capping of long transcripts, whose 5’ ends have moved away from the RNA exit channel, would be less efficient than capping of short transcripts.

Because we found it technically challenging to generate ternary elongation complexes containing long RNA transcripts using DNA:RNA scaffolds, we used our reconstituted enzyme system to compare co-transcriptional capping of short and long transcripts initiated from the Adenovirus major late promoter on the G23 template, which contains two sequential G-less cassettes. Using the approach outlined in Fig. 7A, we first synthesized radiolabeled transcripts of 20 nt and subjected the resulting transcription complexes to high salt washes to remove initiation factors and excess nucleotides. Washed elongation complexes were then extended with unlabeled NTPs to 23mers (short walk) or 223mers (long walk) and incubated with various concentrations of capping enzyme. Since the difference between electrophoretic mobilities of long capped and uncapped transcripts was too small to measure, we included an enzymatic cleavage step post-capping. Reaction products were digested with ribonuclease T1, which cleaves single stranded RNA after G residues, shortening both the 23mers and 223mers to 21 nt, allowing us to easily detect and quantify capping of both short and long transcripts.

**Figure 7.**
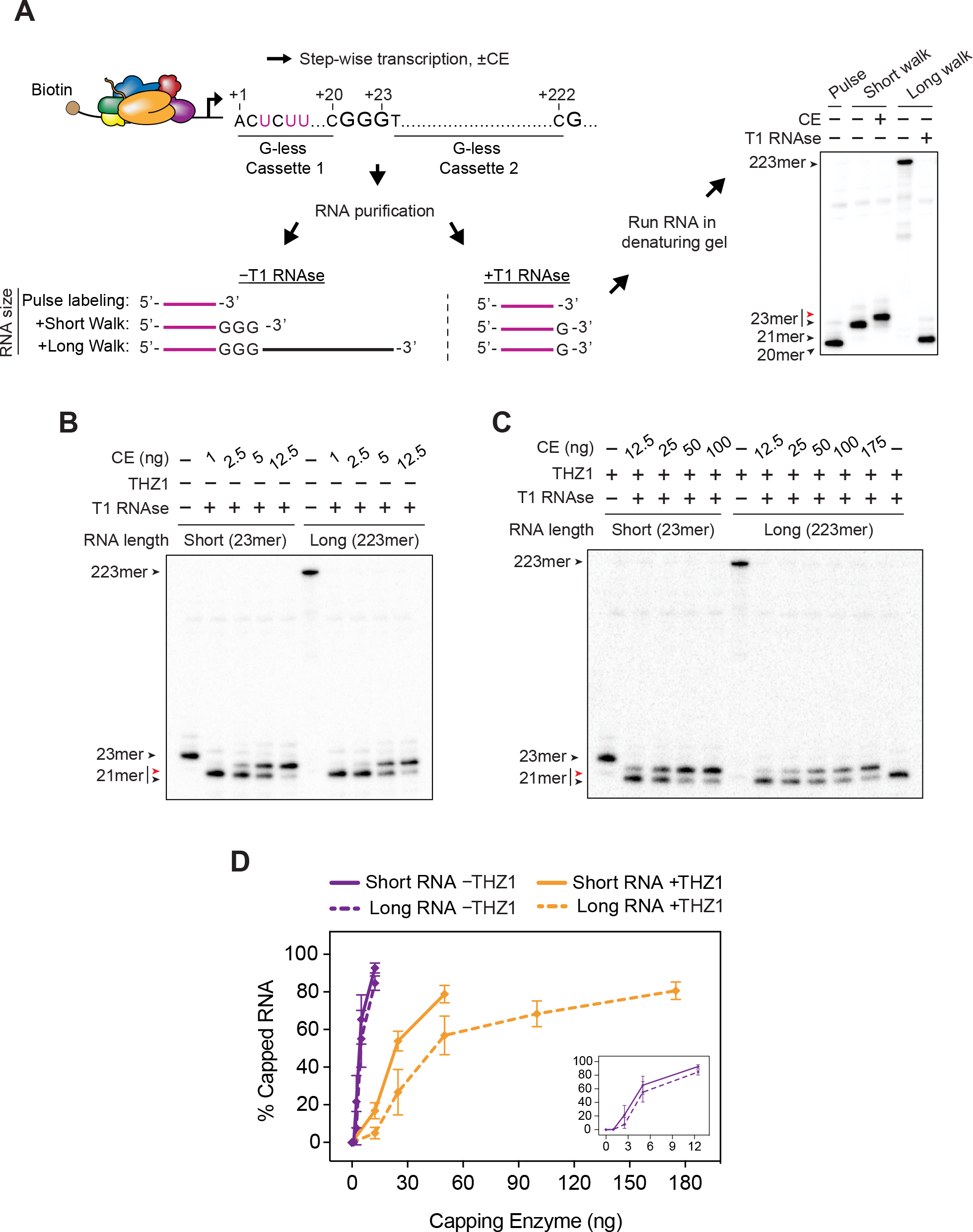
RNA-length dependent co-transcriptional capping. ***A,*** Diagram of method used to assay co-transcriptional capping of short and long RNAs. Transcripts initiated at the AdML promoter on the G23 DNA template were radiolabeled during an initial pulse labeling step and extended to 23mers or 222mers to generate ternary complexes containing short or long RNAs respectively. After incubation with buffer or capping enzyme, short (23-nt) and long RNAs (223-nt) were purified and treated for 15 min with T1 RNAse before denaturing gel electrophoresis. Image is from the same gel as that shown in Fig. 1B. Purple lines and “U” s represent radiolabeled RNAs and nucleotides. Here and in panels ***B*** and ***C***, red and black arrowheads represent capped and uncapped RNA respectively. ***B*** & ***C,*** Ternary complexes containing short (23mer) or long (223mer) RNAs were incubated 50 μM GTP, THZ1 or DMSO, and varying amounts of capping enzyme for 2 min, and reaction products were processed as in ***A***. For the reactions shown in ***C***, 150μM THZ1 was included in all buffers except during washing. ***D,*** The graph shows mean (±S.D.) from at least three independent experiments performed as in panels ***B*** and ***C***. Solid lines, capping of 23mers; dotted lines, capping of 223mers, with (orange) or without (purple) THZ1. Inset shows only reactions without THZ1.

Using this approach, we compared the efficiencies of capping of short and long RNAs, in the presence and absence of the CTD kinase inhibitor THZ1. As shown in Fig. 7D, the efficiency of capping of short and long RNA was indistinguishable in the absence of THZ1, under conditions of maximal CTD phosphorylation. However, in the presence of THZ1 levels that inhibit CTD phosphorylation more than 95%, we observed that increasing the length of nascent RNA reduced the efficiency of capping, although not as dramatically as substituting *S. Pombe* Pol II for mammalian Pol II (an average of 3-fold vs ~10-fold).

The results presented thus far are consistent with the model that the Pol II elongation complex acts as a scaffold that brings capping enzyme and the 5’ end of the nascent transcript together. However, it is formally possible that our observation that free RNA capping cannot be activated in *trans* by phosphorylated Pol II is due not to a requirement that the nascent transcript be tethered to the Pol II elongation complex but rather that passage of the nascent transcript 5’-end through the RNA exit channel during transcript synthesis allows it to adopt a conformation needed for optimal capping. To address this possibility, we investigated the effect on capping of using ribonuclease T1 cleavage to untether the nascent transcript prior to capping.

As diagrammed in figure 8A, we walked Pol II elongation complexes to form a long RNA (Fig. 8B, lane 1). Ternary complexes were then washed and incubated with T1 RNAse before capping to generate shorter RNA fragments that have passed through the RNA exit channel during synthesis but are untethered from the elongation complex, or, in control reactions, after capping. Consistent with results shown earlier, 5 ng of capping enzyme was sufficient to cap about 50% of nascent transcripts tethered to ternary elongation complexes (Fig. 8B, compare lanes 2 and 3). In contrast, when nascent transcripts were untethered from elongation complexes by treatment with T1 before addition of capping enzyme, capping efficiency was dramatically reduced even at higher concentrations of capping enzyme (175 ng), resembling the capping kinetics of free RNA added in *trans* (Figure 8B). Thus, untethering nascent RNA from the Pol II elongation complex was sufficient to render the efficiency of its capping similar to that observed with free RNA, providing further support for the tethering model for activation of co-transcriptional capping.

**Figure 8.**
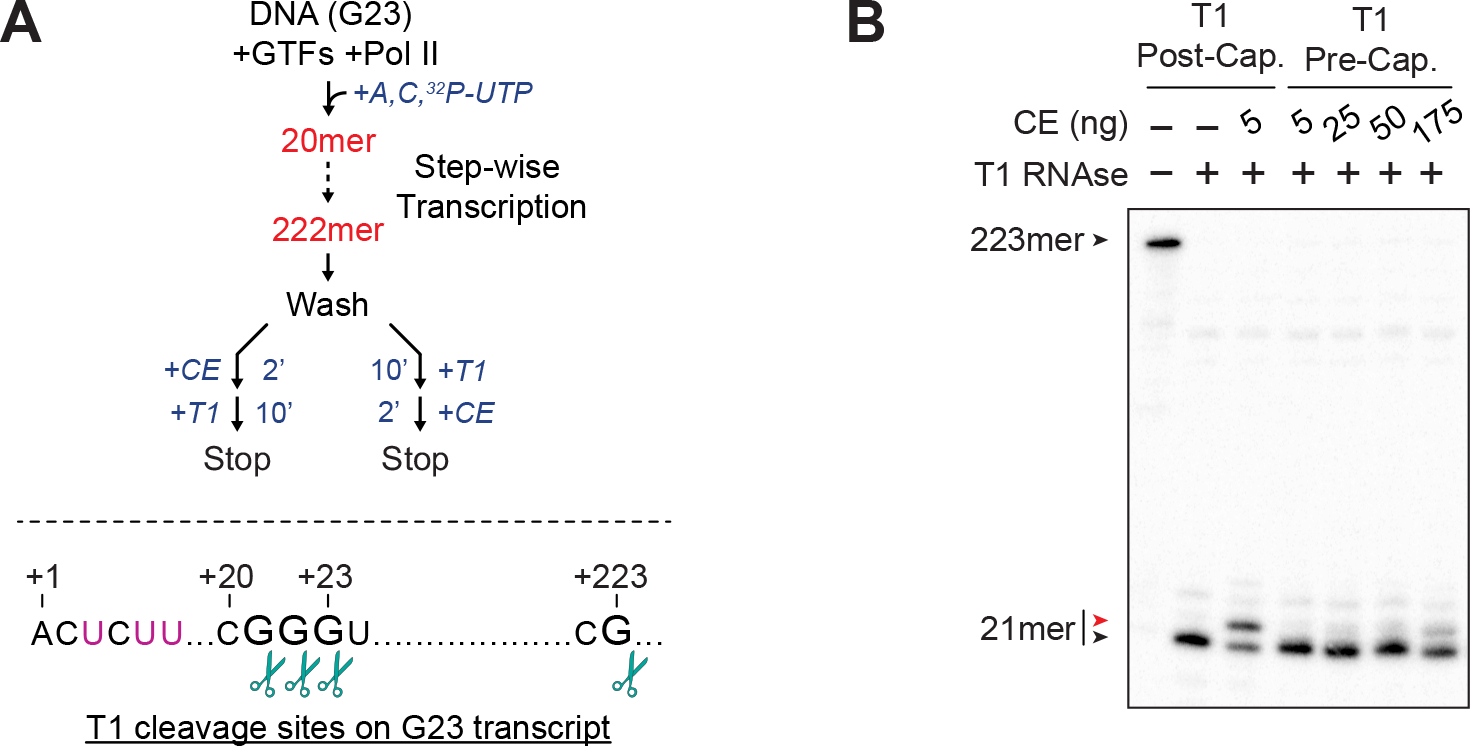
Release of nascent RNA from ternary complex prevents capping activation. ***A,*** (Top) Diagram of reaction scheme. Ternary complexes initiated from the AdML promoter were walked to position +222, washed, resuspended in buffer with 50 μM GTP, and treated with T1 RNase for 10 min post-or pre-capping. When T1 was added after capping, reactions were incubated with 5 ng of capping enzyme for 2 min, then incubated with T1 RNase and 10 mM EDTA for 10 min. When T1 was added after capping, reactions were incubated with T1 RNase for 10 min, then incubated with the indicated amounts of capping enzyme for 2 min. (Bottom) Diagram of the RNA sequence associated with ternary complexes, showing T1 cleavage sites (scissors). Purple indicates radiolabeled nucleotides. ***B,*** Analysis of the capping reactions prepared in ***A***. Red and black arrowheads represent capped and uncapped RNA respectively.

## Discussion

In this report, we investigate the biochemical mechanism underlying co-transcriptional capping activation of mammalian Pol II transcripts. Our findings are consistent with the model that activation of co-transcriptional capping is primarily due to tethering both the nascent transcript and the capping enzyme to transcribing Pol II *via* contacts with Ser5-phosphorylated CTD and yet-to-be-defined sites on the body of Pol II.

First, we observe robust activation of capping of nascent transcripts in reactions containing purified Pol II assembled into artificial ternary complexes. Co-transcriptional capping of transcripts in these artificial ternary complexes is enhanced about 10-fold in the presence of catalytically active TFIIH kinase. We note that blocking TFIIH kinase activity with THZ1 in promoter-dependent assays leads to an ~5-fold decrease in capping efficiency. We expect this difference is due to residual CTD kinase activity in reactions containing THZ1, since as shown in Fig. 2A THZ1 greatly reduces, but does not completely inhibit, CTD phosphorylation. TFIIH-dependent capping activation is not observed in reactions containing mutant Pol II lacking CTD, providing strong support for the notion that the Pol II CTD is the sole target for phosphorylation-dependent activation of co-transcriptional capping in this system.

Second, our observation that the Pol II CTD contributes to activation of capping only in the presence of TFIIH and under conditions where the TFIIH CDK7 kinase phosphorylates the CTD is consistent with a large body of prior studies arguing that the phosphorylated Pol II CTD plays an important role in activation of co-transcriptional capping (Cho et al., 1998; Cho et al., 1997; Ho and Shuman, 1999; Ho et al., 1998; McCracken et al., 1997; Rodriguez et al., 2000). Consistent with these early observations, a recent study demonstrated that CTD Ser5 phosphorylation and capping enzyme occupancy at the 5’ end of genes is reduced genome-wide in human cells expressing an analog-sensitive CDK7 mutant (Ebmeier et al., 2017). These studies brought to light two potential roles for the phosphorylated CTD in activation of capping. First, capping enzyme is recruited to the Pol II elongation complex *via* specific and stable binding to the phosphorylated CTD. In addition, capping enzyme can be allosterically activated to form the capping enzyme-GMP intermediate upon interaction with serine 5 phosphorylated CTD peptides. Based on our evidence (i) that addition in *trans* of Pol II elongation complexes, with or without TFIIH, had no effect on efficiency of free RNA capping, (ii) that ribonucleolytic release of nascent transcripts from Pol II elongation complexes abolished activation of capping, and (iii) that CTD phosphorylation has a much greater effect on the rate of capping than on formation of the capping enzyme-GMP intermediate, we argue that the critical role of the phosphorylated CTD in activation of co-transcriptional capping is most likely recruitment and tethering of the capping enzyme to Pol II.

Third, our observation that robust activation of co-transcriptional capping occurs even in the absence of the Pol II CTD or TFIIH supports the model that there is a parallel CTD-independent mechanism for activation of capping. That Pol II from the fission yeast *S. pombe* fails to support robust activation of co-transcriptional capping by mammalian capping enzyme unless the CTD is phosphorylated is consistent with the ideas (i) that species-specific interactions of mammalian capping enzyme with mammalian Pol II are critical for this CTD-independent mechanism for activation of capping and (ii) that tethering of capping enzyme to Pol II is governed by a multipartite Pol II binding site, which includes the phosphorylated CTD and a site(s) outside the CTD. Notably, phosphorylation of the fission yeast Pol II CTD is sufficient to restore activation of co-transcriptional capping by mammalian capping enzyme to levels seen with mammalian Pol II, illustrating that the CTD-dependent and CTD-independent pathways for activation of capping function in parallel and can compensate for each other under some conditions. In light of these findings, it is noteworthy that Schwer & Shuman reported that in fission yeast, an otherwise lethal mutation of Rpb1—mutation of Serine 5 of all CTD repeats to alanine—can be rescued with mammalian capping enzyme, but only when capping enzyme is covalently tethered to the CTD mutant (Schwer and Shuman, 2011). Thus, even in the absence of species-specific interactions between capping enzyme and the Pol II body, the forced proximity of capping enzyme to the ternary complex, brought together by the covalent tether, is sufficient to promote efficient co-transcriptional RNA capping and, therefore, survival.

A similar model has been proposed for budding yeast, where the most stable physical interaction between capping enzyme and Pol II required two interfaces on Pol II: CTD phosphorylated on serine 5 and the foot domain on Rpb1 (Suh et al., 2010). Disruption of either interaction caused severe growth defects in vivo and interfered with physical association of capping enzyme with Pol II. Yet, recent cryo-EM analysis of a ternary complex bound to the capping machinery was not able to confirm this interaction with the foot domain, but instead concluded that capping enzyme spanned the end of the Pol II RNA exit tunnel, where it would be positioned to capture the nascent transcript as it emerges from polymerase (Martinez-Rucobo et al., 2015). The degree to which either of these interactions contribute directly to co-transcriptional capping in yeast or whether additional, yet to be defined contacts are required remains to be determined. In any case, our observation that long transcripts are capped as efficiently as short transcripts that have just emerged from the exit tunnel when the CTD is phosphorylated argues against a model in which optimal co-transcriptional capping requires that capping enzyme must capture the 5’end of the nascent transcript as it emerges from Pol II.

Finally, our evidence that co-transcriptional capping in artificial elongation complexes can be strongly activated by parallel pathways involving contacts with phosphorylated CTD and with the surface of Pol II may provide insight into recent findings from Nilson and colleagues (Nilson et al., 2015). They observed that capping of RNA in transcription complexes that had been assembled in nuclear extracts and washed with high salt was much less sensitive to inhibition of CTD phosphorylation with THZ1 than was capping in low salt washed complexes. Based on this observation, they proposed that the major function of CDK7 kinase in capping regulation is to promote dissociation of an activity that interferes with capping. While the identity of such an activity remains unknown, our results are consistent with the model that a factor(s) bound to the body of Pol II could occlude binding site(s) for capping enzyme and thereby interfere with capping when the CTD is not phosphorylated. We believe, however, that it is not necessary to postulate that CDK7-dependent phosphorylation events are required to remove such a factor from the transcription complex. As shown by the results of our experiments using mammalian capping enzyme with *S. pombe* ternary complexes, phosphorylation of the CTD could lead to strong activation of co-transcriptional capping by compensating for the lack of contacts between capping enzyme and sites on the Pol II body rendered inaccessible by binding of the proposed factor(s) to Pol II. In the future, it will be of considerable interest to explore these issues in more detail.

## Acknowledgements

We thank S. Shuman for providing the mammalian capping enzyme cDNA, Henrik Spahr for *S. pombe* Pol II, and D. Wang and J. M. Egly for helpful discussions. This work was supported in part by a grant to the Stowers Institute for Medical Research from the Helen Nelson Medical Research Fund at the Greater Kansas City Community Foundation.

## Author contributions

M.N.G., J.W.C., and R.C.C. designed research, M.N.G. performed research, C.T.-S. and S.S. provided key reagents, and M.N.G., J.W.C., and R.C.C. wrote the paper.

---

**Supplementary Figure 1.**
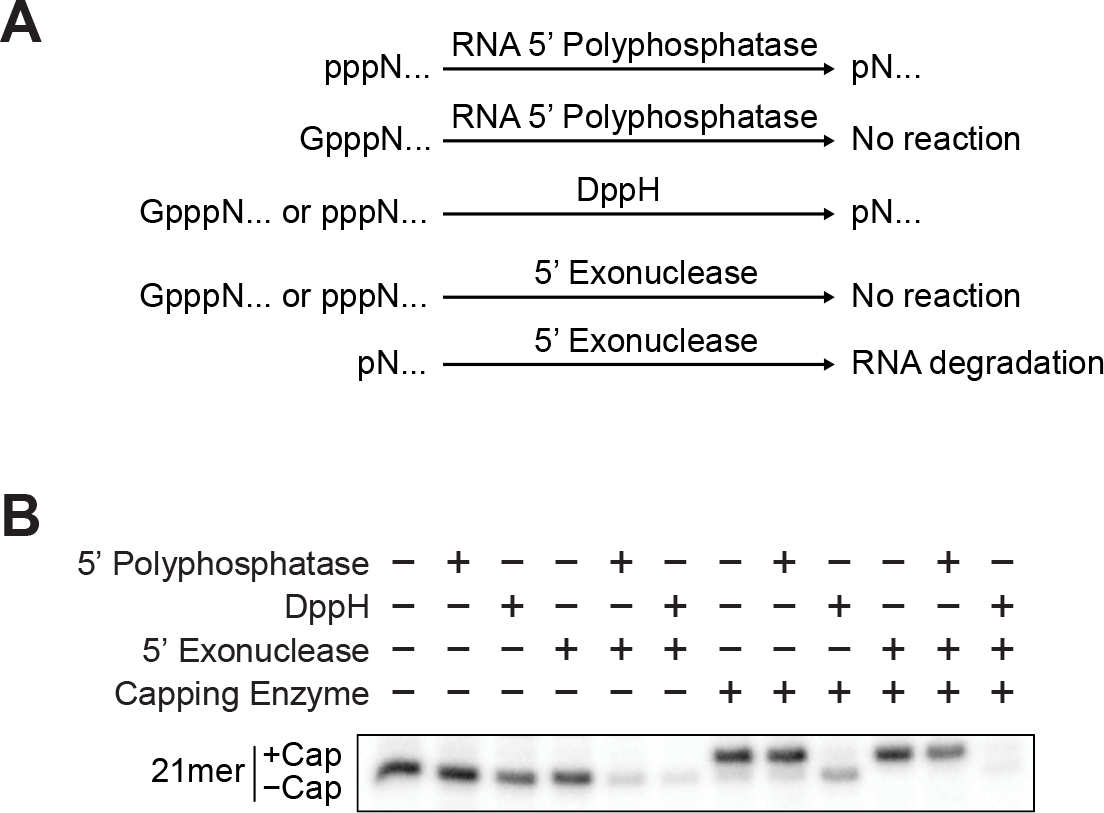
Enzymatic cleavage of capped RNA. ***A,*** Diagrams show products of reactions catalyzed by each enzyme in the presence of capped (GpppN…) or uncapped (pppN…) RNAs. Equations show catalytic reactions of each enzyme when mixed with capped or uncapped transcripts. RNA 5’ polyphosphatase converts 5’ triphosphate (pppN) to 5’ monophosphate (pN) only when RNA is uncapped; Decapping pyrophosphohydrolase (Dpph) converts 5’ triphosphate to 5’ monophosphate when RNA 5’ ends ore capped or uncapped; 5’ exonuclease degrades only RNA having 5’ monophosphate ends. ***B,*** Synthesis and capping of 21mers associated with transcription complexes initiated from the AdML was performed in reactions containing 100 μM 3’OMeG and 100 ng of capping enzyme for 30 min, and RNA was purified as described in Materials and Methods. Dephosphorylation reactions were carried out for 1 hr at 30 °C in 20 μl reaction mixtures containing purified radiolabeled RNA, 2μl of the appropriate 10 x enzyme buffer provided with each enzyme, with or without 2 μl of 5’ RNA polyphosphatase (Epicentre) or 2 μl of DppH (Tebu-bio). RNA was purified again and incubated for 1 hr at 30 °C in 20 μl reactions containing 2 μl 10x exonuclease buffer, with or without 2 μl of 5’ exonuclease.

**Supplementary Figure 2.**
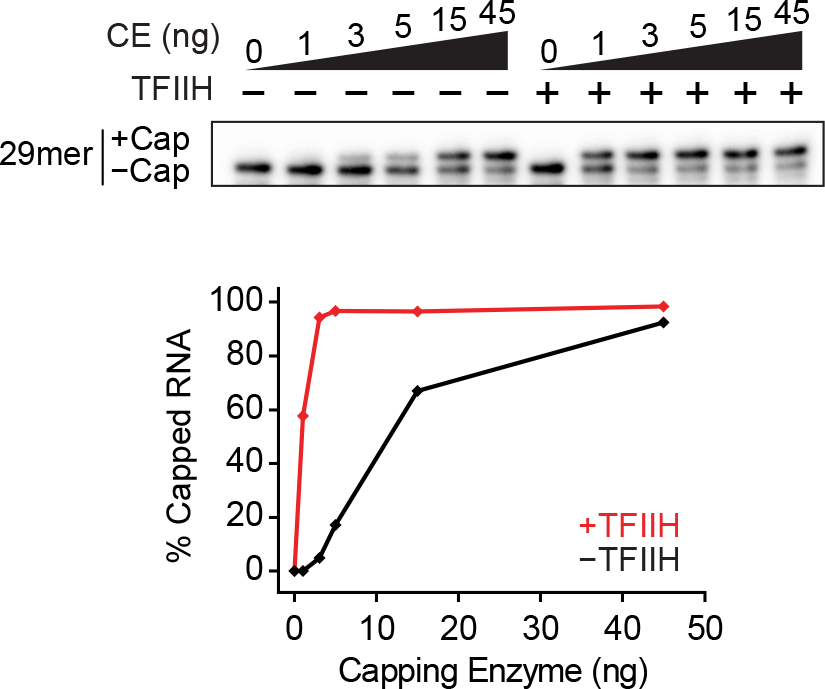
Titration of capping enzyme shows TFIIH-dependent activation of capping in artificial ternary complexes. Artificial ternary complexes containing 29mers were incubated with or without 300 ng of purified TFIIH for 10 minutes, washed, and resuspended in buffer with 50 μM GTP and the indicated amounts of capping enzyme. Capping reactions were stopped after 15 min. The graph shows quantification of RNA capping in these reactions.

**Supplementary Figure 3.**
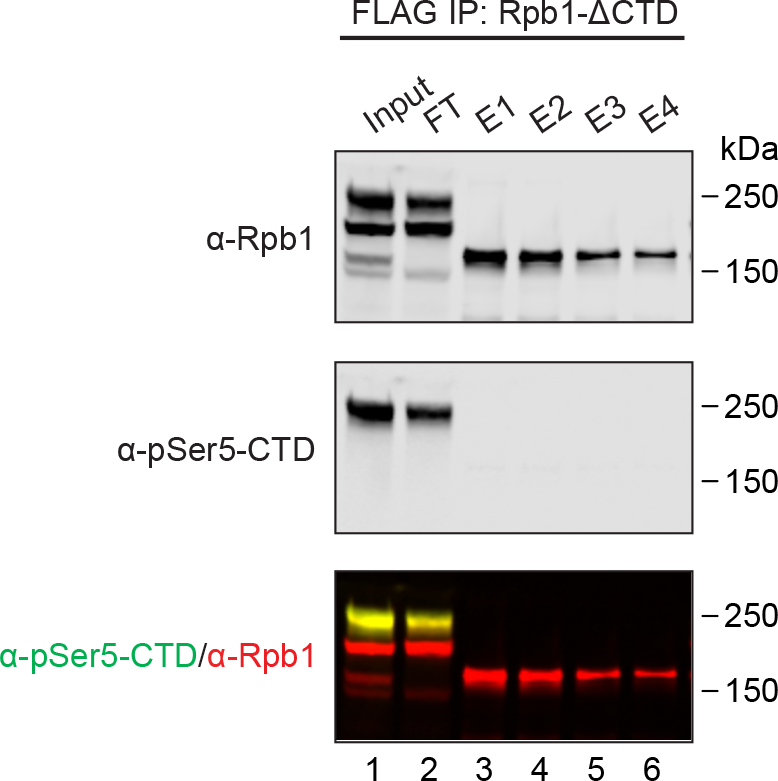
Purification of Pol II lacking the CTD from stable cell lines. Western blot showing endogenous Rpb1 and F:Rpb1-ΔCTD through the stages of flag immunopurification using antibodies against total Rpb1 (α-Rpb1) or Ser5 phosphorylated Rpb1 CTD (α-pSer5-CTD). See “Materials & Methods” for details. FT, FLAG agarose flow through fraction; E1 through E4, fractions obtained during elution with FLAG peptide from anti-FLAG agarose.

